# Toward a model-free feedback control synthesis for treating acute inflammation

**DOI:** 10.1101/294389

**Authors:** Ouassim Bara, Michel Fliess, Cédric Join, Judy Day, Seddik M. Djouadi

## Abstract

An effective and patient-specific feedback control synthesis for inflammation resolution is still an ongoing research area. A strategy consisting of manipulating a pro and anti-inflammatory mediator is considered here as used in some promising model-based control studies. These earlier studies, unfortunately, suffer from the difficultly of calibration due to the heterogeneity of individual patient responses even under similar initial conditions. We exploit a new model-free control approach and its corresponding “intelligent” controllers for this biomedical problem. A crucial feature of the proposed control problem is as follows: the two most important outputs which must be driven to their respective desired states are sensorless. This difficulty is overcome by assigning suitable reference trajectories to the other two outputs that do have sensors. A mathematical model, via a system of ordinary differential equations, is nevertheless employed as a “virtual” patient for *in silico* testing. We display several simulation results with respect to the most varied situations, which highlight the effectiveness of our viewpoint.

## 1 Introduction

Inflammation is a key biomedical subject (see, *e.g.*, [58,86]) with fascinating connections to diseases like cancer (see, *e.g*., [10]), AIDS (see, *e.g*., [27]) and psychiatry (see, *e.g*., [56]). Mathematical and computational models investigating these biological systems have provided a greater understanding of the dynamics and key mechanisms of these processes (see, *e.g*., [85]). The very content of this particular study leads citations of mathematical models where differential equations play a prominent role (see, *e.g*., [5,6,7,18,23,24,26,28,30,31,40,46,51,59,61,63,64,65,75,83,92]). The usefulness of those equations for simulation, prediction purposes, and, more generally, for understanding the intimate mechanisms is indisputable. In addition, some models have also been used in order to provide a real-time *feedback control* synthesis (see, *e.g*., [8] for an excellent introduction to this important engineering topic) for treating acute inflammation due to severe infection. Insightful results were obtained via two main model-based approaches:

- optimal control [12,14,15,44,76,77,78,80],
- model predictive control [25,41,69,96].

Our work in [12,14,15,25,96] made use of the low dimensional system of ordinary differential equations (ODE) derived in [65] (see also [26]). This four variable model possesses the following characteristics:

- The model is based on biological first principles, the non-specific mechanisms of the innate immune response to a generic gram-negative bacterial pathogen.
- A variable representing anti-inflammatory mediators e.g. Interleukin-10, Transforming Growth Factor-*β*) is included and plays an important role in mitigating the negative effects of inflammation to avoid excessive tissue damage.
- Though a qualitative model of acute inflammation, it reproduces several clinically relevant outcomes: a healthy resolution and two death outcomes.

The calibration of a system of differential equations can be quite difficult since the identification of various rate parameters requires specific data in sufficient quantities, which may not be feasible. Additionally, there is much heterogeneity to account for between patient responses such as the initiating circumstances, patient co-morbidities and personal characteristics, like genetics, age, gender, In spite of promising preliminary results in [11,13,96], state estimation and parameter identification of highly nonlinear models may still require more data than can be reasonably collected. These roadblocks hamper the use of model-based control strategies in clinical practice in spite of recent mathematical advances. Here, another route, *i.e*, *model-free control (MFC)* and the corresponding “intelligent” feedback controllers [34], are therefore explored.^1^ We briefly introduce the method before discussing its application to the scenario of controlling the inflammatory response to pathogen. We begin by replacing the poorly known global description by the *ultra-local model* given by:

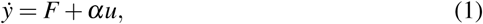

where

- the control and output variables are *u* and *y*, respectively;
- the derivation order of *y* is 1 like in most concrete situations;
- *α* ∈ ℝ is chosen by the practitioner such that *αu* and *ẏ* are of the same magnitude;
- *F* is estimated via the measurements of *u* and *y*;
- *F* subsumes not only the unknown system structure but also any perturbation.

*Remark 1* The following comparison with computer graphics is borrowed from [34]. To produce an image of a complex curve in space, the equations defining that curve are not actually used but, instead an approximation of the curve is made with short straight line segments. Equation (1), which might be viewed as an analogue of such a segment, should hence not be considered as a global description but instead as a rough linear approximation.

*Remark 2* The estimation of the fundamental quantity *F* in Equation (1) via the control and output variables *u* and *y* will be detailed in Section 2.2. It connects our approach to the *data-driven* viewpoint which has been adopted in control engineering (see, *e.g.*, [39,42,67,68]) and in studies about inflammation (see, *e.g.*, [9,20,69,84]).

Ideally, data associated with the time courses of the inflammatory response variables would be generated by measurements from real patients. In our case, it would be the pro-inflammatory and anti-inflammatory variables of the model (patient) which we would want to track; and therefore, define these as the reference trajectories (available data) for the model-free setup. Once the quantity *F_est_* is obtained, the loop is closed by an *intelligent proportional controller*, or *iP*:

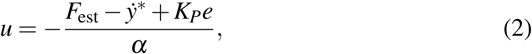

where

- *F_est_* is an estimate of *F*;
- *y*\* is the reference trajectory;
- *e* = *y-y*\* is the tracking error; and
- *K_P_* is an usual *tuning gain.*

With a “good” estimate *F_est_* of *F*, *i.e*., *F* – *F_est_* ≃ 0, Equations (1)-(2) yield

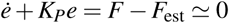

Thus *e*(*t*) ≃ *e*(0) exp(—*K_P_t*), which implies that

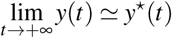

if and only if, *K_P_* > 0. In other words, the scheme ensures an excellent tracking of the reference trajectory. This tracking is moreover quite robust with respect to uncertainties and disturbances which can be numerous in a medical setting such as considered here. This robustness feature is explained by the fact that *F* in Equation (1) encompasses “everything,” without trying to distinguish between its different components. In our application, sensorless outputs must be driven in order to correct dysfunctional immune responses of the patient. Here, this difficult problem is solved by assigning suitable reference trajectories to those systems variables which can be measured. This *feedforward* viewpoint is borrowed from the *flatness-based* control setting [37] (see also [8,48,73]).

After justifying model-free control in Section 2, Section 3 presents results from applying the method to a heterogeneous *in silico* virtual patient population generated in [25]. The cohort of virtual patients are summarized in Section 3.1. The computer simulations demonstrate the great robustness of the model-free control strategy with respect to noise corruption, as demonstrated in Section 3.4. Concluding remarks in Section 4 discuss some of the potential as well as the remaining challenges of the approach in the setting of controlling complex immune responses.

A first draft has already been presented in [16].

## 2 Justification of the model-free approach: A brief sketch

### 2.0.1 Justification of the ultra-local model

We first justify the ultra-local model given in (1). For notational simplicity, we restrict to a system with a single control variable *u* and a single output variable *y*. Assume that the system is a *causal*, or *non-anticipative functional*; In other words, for any time instant *t* > 0, let

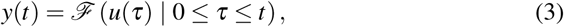

where *ℱ* depends on

- the past and present but not the future,
- various perturbations, and
- initial conditions at *t* = 0.

*Example 1* A representation of rather general nonlinear functionals, also popular in the biological sciences, is provided by a Volterra series (see, *e.g*., [45]):

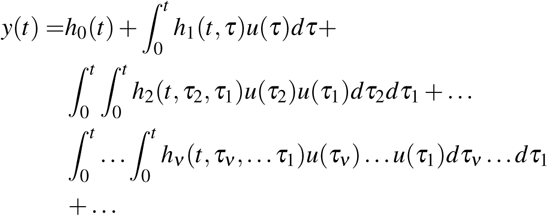

Solutions of quite arbitrary ordinary differential equations, related to input-output behaviors, may be expressed as a Volterra series (see, *e.g.*, [36]).

Let

- *ℐ* ⊂[0, +∞[be a compact subset and
- 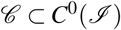 be a compact subset, where *C*^0^(*ℐ*) is the space of continuous functions *ℐ* → ℝ, which is equipped with the topology of uniform convergence.

Consider the Banach ℝ-algebra 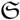 of continuous causal functional 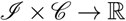 If a subalgebra contains a non-zero constant element and separates points in 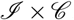 then it is dense in 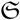 according to the classic Stone-Weierstraß theorem (see, *e.g.*, [70]). Let 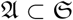 be the set of functionals which satisfy an algebraic differential equation of the form

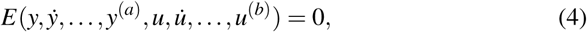

where *E* is a polynomial function of its arguments with real coefficients. It has been proven in [34] that with this, the conditions of the Stone-Weierstraß theorem are satisfied and, therefore, 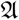 is dense in 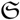.

Assume therefore that our system is “well” approximated by a system defined by Equation (4). Let v be an integer, 1 ≤ *v* ≤ *a*, such that

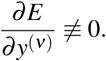

The implicit function theorem yields locally

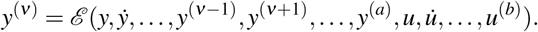

This may be rewritten as

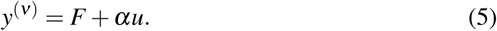

In most concrete situations such as the one here, the order *v* = 1 of derivation, as in Equation(1), is enough. See [34] for an explanation and for some examples where *v* = 2.

### 2.1 Closing the loop

If *v* = 1 in Equation (5), we are back to Equation (1). The loop is closed with the intelligent proportional controller (2).

### 2.2 Estimation of *F*

Any rather general function [*a, b*] → ℝ *a*, *b* ∈ ℝ, *a* < *b*, may be approximated by a *step* function *F*_approx_, *i.e.*, a piecewise constant function (see, *e.g*., [71]). Therefore, for estimating a suitable approximation of *F* in Equation (5), the question reduces to the identification of the constant parameter *Ф* in

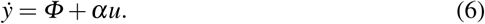

Here a recent real-time algebraic estimation/identification techniques are employed ([38,74]). With respect to the well-known notations of operational calculus (see, *e.g*., [32,93]) (which are identical to those of the classic Laplace transform taught everywhere in engineering *e.g*., [29], [8]), Equation (6) yields:

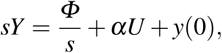

where *U* and *Y* are the operational analogues of *u* and *y.* In the literature, *U* and *Y* are often called the *Laplace transforms* of *u* and *y*, and s is the *Laplace variable* (see, *e.g.*, [8]).

We eliminate the initial condition y(0) by left-multiplying both sides by 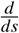 or, in other words, by differentiating both sides with respect to s:

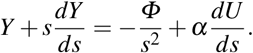

The product by *s* corresponds in the time domain to the derivation with respect to time. Such a derivation is known to be most sensitive to noise corruptions. Therefore, multiply both sides on the left by *s*^-2^ in order to replace derivations by integrations with respect to time, which are quite robust with respect to noise (see [33] for more explanations). Recall that 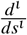, where *ι* ≥ 1 is an integer, corresponds in the time domain to the multiplication by (*-t*)^*ι*^. Then

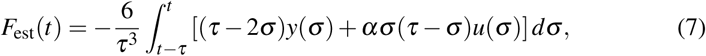

where *τ* > 0 might be quite small. This integral may of course be replaced in practice by a classic digital filter.

There are other formulas one can use for obtaining an estimate of *F*. For instance, closing the loop with the iP (2) yields:

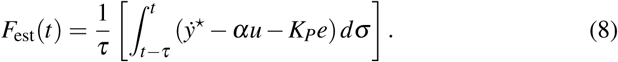

*Remark 3* Measurement devices are always corrupted by various noise sources (see, *e.g*., [79]). The noise is usually described via probabilistic/statistical laws that are difficult to write down in most concrete situations. Following [33] where *nonstandard analysis* is used, the noise is related to quick fluctuations around zero [22]. Such a fluctuation is a Lebesgue-integrable real-valued time function *ℱ* which is characterized by the following property: the integral of *ℱ* over any finite time interval, 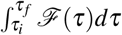, is infinitesimal. Therefore, noise is attenuated thanks to the integrals in formulas (7)-(8).

## 3 Computer Simulation

### 3.1 Virtual patients

In [25], a cohort of virtual patients were defined by using the ODE model of [65] for the underlying immune response dynamics for each patient and with each differing in the value of six of the rate parameters and two of the initial conditions. This same cohort was used in this study as well. The ODE model describes an abstract dynamical representation of an acute inflammatory response to pathogenic infection:

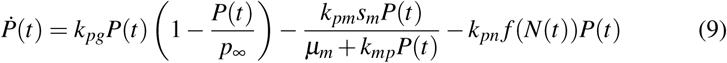

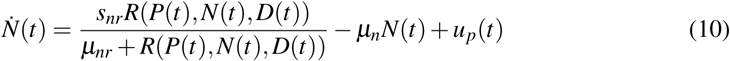

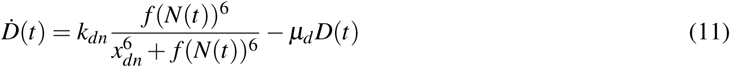

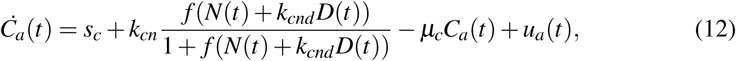

where

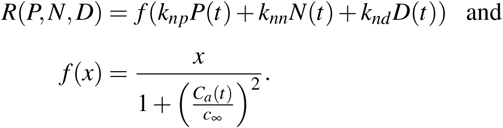

- Equation (9) represents the evolution of the bacterial pathogen population *P* that causes the inflammation.
- Equation (10) governs the dynamics of the concentration of a collection of early pro-inflammatory mediators such as activated phagocytes and the pro-inflammatory cytokines produced by *N*.
- Equation (11) corresponds to tissue damage (D), which helps to determine response outcomes.
- Equation (12) describes the evolution of the concentration of a collection of anti-inflammatory mediators *C_a_*.

As explained in [65], *f*(*x*) represents a Hill function that models the impact of activated phagocytes and their by-products (*N*) on the creation of damaged tissue. With this modeling construct, tissue damage (*D*) increases in a switch-like sigmoidal fashion as *N* increases such that it takes sufficiently high levels of *N* to incite a moderate increase in damage and that the increase in damage saturates with sufficiently elevated and sustained *N* levels. The hill coefficient (exponent) 6 was chosen to model this aspect which also ensured that the healthy equilibrium had a reasonable basin of attraction for the *N/D* subsystem.

For the reference set of parameter values which is given in Table I of [65], the above model possesses three (positive) stable equilibria which can be qualitatively interpreted as the following clinical outcomes:

- Healthy outcome: equilibrium in which *P* =*N* =*D* = 0 and *C_a_* is at a background level.
- Aseptic death outcome: equilibrium in which all mediators *N*, *C_a_*, and *D* are at elevated levels, while pathogen, *P*, has been eliminated.
- Septic death outcome: equilibrium in which all mediators *N, C_a_*, and D together with the pathogen P are at elevated levels (higher than in the aseptic death equilibrium).

Note that the model was formulated to represent a most abstract form of the complex processes involved in the acute inflammatory response. Hence, as explained in [65] the variables *N* and *C_a_* represent multiple mediators with similar inflammatory characteristics, and *D* is an abstract representation of collateral tissue damage caused by inflammatory by-products. This abstraction reduces the description to four essential variables which also allows for tractable mathematical analysis. Therefore, the units of these variables are in arbitrary units of *N*-units, *C_a_*-units, *D*-units, since they represent various types of cells and thus, they qualitatively, rather than quantitatively, describe the response of the inflammatory mediators and their by-products. Pathogen, *P*, units are more closely related to numbers of pathogens or colony forming units (CFU), but abstract units *P*-units are simply used as well and this population is scaled by 10^6^/cc. More details about the model development can be found in [65].

The diagram in Figure 1 characterizes the different interactions between the states of the inflammatory model. A solid line with an arrow head indicates an up-regulation, whereas a dashed line with circular head indicates inhibition or down-regulation of a process. For instance, early pro-inflammatory mediators, *N*, respond to the presence of pathogen, *P*, by initiating self-recruitment of additional inflammatory mediators and *N* is therefore up-regulated by the interaction with P to attempt to efficiently eliminate the pathogen. The self up-regulation that exists for *P* is due to replication. Furthermore, *N* inhibits *P* by eliminating it at some rate. The inflammation caused by *N*, however, results in tissue damage, D, which can provide a positive feedback into the early inflammatory mediators depending on their intensity. To balance this, anti-inflammatory mediators, such as cortisol, IL-10, and TGF-*β*, can mitigate the inflammation and its harmful effect by suppressing the response by *N* and the effects of D in various ways. The *C_a_* variable maintains a small positive background level at equilibrium in the presence of no pathogen. Following the setup used in [25], the reference parameter value for *C_a_*(0) is set to 0.125 and virtual patients have a value that is ±25*%* of the reference value. In addition, the values of six other parameters as well as the initial condition for P are set to have differing (positive) values from the reference set. In particular, the values of these parameters and initial conditions were generated from a uniform distribution on defined parameter ranges or on a range that was +/-25% of the (mean) reference value. The remaining parameters retained the same values as those in the reference set. These differences distinguish one virtual patient from another.

**Fig. 1:**
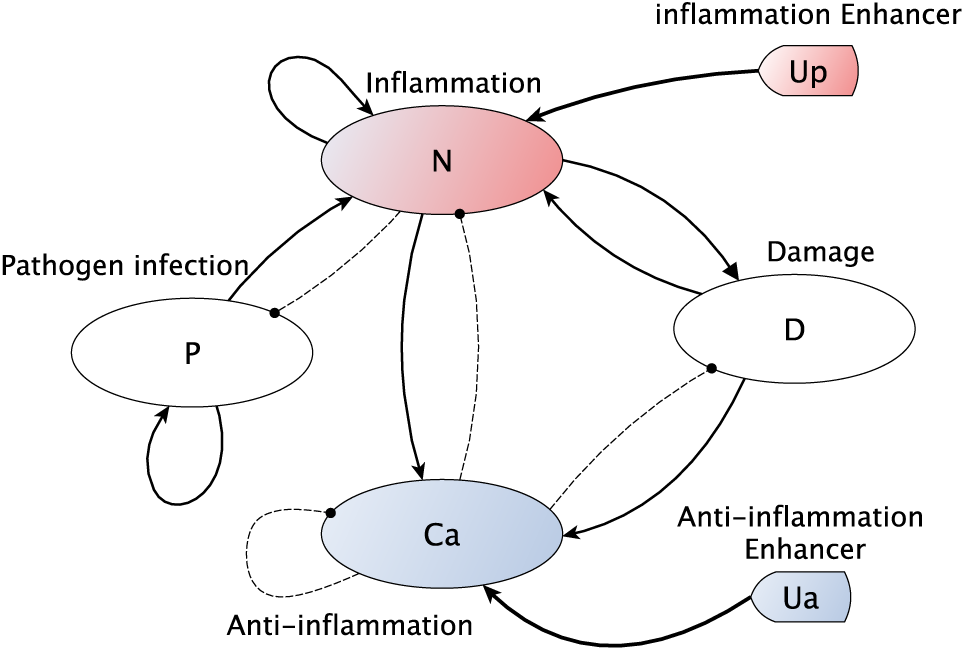
Diagram of the mediators of the acute inflammatory response to pathogen as abstractly modeled in [65]. Solid lines with arrow heads and dashed lines with nodes/circular heads represent upregulation and inhibition, respectively. *P*: replicating pathogen, *N*: early pro-inflammatory immune mediators, *D*: marker of tissue damage/dysfunction caused by inflammatory response, *C_a_*: inhibitory anti-inflammatory mediators, *u_a_* and *u_p_*: time-varying input controls for the anti- and pro-inflammatory therapy, respectively.

We use the set of 1000 virtual patients generated by [25] in the way described above to evaluate the performance of the proposed control strategy. The set of patients was classified with respect to their outcome after an open loop simulation for a long enough time to numerically determine outcome without ambiguity. Of the 1000 virtual patients, 369 did not resolve the infection and/or inflammatory response on their own and succumbed to a septic (141) or aseptic (228) death death outcome. On the other hand, 631 exhibited a healthy outcome of which there were two distinct subsets:

1. - 379 of the 631 healthy virtual patients did not necessitate treatment intervention because their inflammatory levels did not exceed a specified threshold (defined to be *N*(*t*) ≤ 0.05, set in [25]). These virtual patients were excluded from receiving treatment and from our *in silico* study.
2. - The remaining 252 of these virtual patients did surpass the specified threshold, *N*(*t*) ≥ 0.05, and are included in the cohort that receives treatment. However, these virtual patients would be able to resolve to health on their own, in the absence of treatment intervention. An important issue for these particular virtual patients is not to harm them with treatment.

Thus, 621 of the 1000 generated virtual patients receive treatment via our control design. Once a suitable reference trajectory is provided for the states with sensors, the derivation of the control part is straightforward which we now discuss.

### 3.2 Control design

As in previous control studies using this model, we assume that the state components *P* and *D* in Equations (9) and (12) are not measurable; whereas, the states *N* and *C_a_*in Equations (10) and (12), respectively, are:

- easily measured and
- influenced by the control variables *u_p_* and *u_a_*, respectively.

We then introduce two equations of type (1):

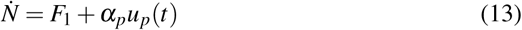

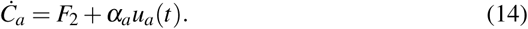

We emphasize that, like in [47], the above two ultra-local systems may be “decoupled” so that they can be considered as monovariable systems. It should be nevertheless clear from a purely mathematical standpoint that *F*_1_ (resp. *F*_2_) is not necessarily independent of *u_a_* (resp. *u_p_*). The two corresponding iPs (2) then read

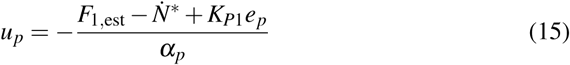

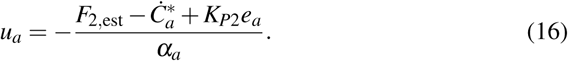

The tracking errors are defined by

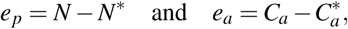

where *N*\* and 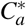 represent the reference trajectories corresponding to pro and anti-inflammatory measurements *N* and *C_a_*, respectively. Knowing that *F* encapsulates all the model uncertainties and disturbances as already explained in the introduction, a good estimate *F_est_* provides exponential local stability of the closed-loop system. The following algorithm 1 provides a good summary on the functioning of the proposed methodology for immune regulation:

#### Algorithm 1 Model-free Control

**Step 1: Initialization**, k = 0

*u_p_*(*0*) = 0, define reference trajectories *N*\*, initialize *K_p_* and *α*, fix the sampling time *T_e_*;

For 1 ≤ *k* ≤*T_f_*

**Step 2**,: Get measurements of *N* and *u_p_*;

**Step 3** Estimation of *F*:

Estimate *F* according to a discrete implementation of equation (7);

**Step 4**: Close the loop according to equation (15) and return to **Step 2**.

Note that the same design procedure can lead to the derivation of the control *u*_a_; however, this time we associate the measurement *C_a_* with the control *u_a_* (See equation (14)). The interesting fact about this approach is that we do not need to control the state variables *P* and *D*, which are not measurable. Solving the tracking problem consisting of following closely the reference trajectory of *N* and *C_a_* is enough to drive the pathogen and damage to values in the basin of attraction of the healthy equilibrium where they would converge to this state as time progressed, thereby ‘curing’ the patient.

### 3.3 Results without noise corruption

We first examine the performance of the control approach with respect to the set of virtual patients and their individual corresponding initial conditions. The robustness of the control law with the addition of corrupting measurement noise will be discussed afterward. In what follows, the reference trajectories which are inspired from [15], correspond to the measurable states *N* and *C_a_*. They will be highlighted in dashed lines.

The simulations for all patients were performed under the following conditions:

- a sampling time of 1 minute,
- *α_p_* = 1, *α_a_ =* 10 in Equations (13)-(14),
- *K_P_*_1_ = *K_P_*_2_ = 0.5 in Equations (15)-(16), and
- 250 hours simulation time to numerically determine outcomes without ambiguity; though we note stress that our control objectives were reached in less than 250 hours.

The use of the same reference trajectory for all simulations emphasizes the robustness of the proposed control approach with respect to the variability among virtual patient parameter values and initial conditions. Figure 2 represents a successful outcomes related to 92 out of the 141 septic patients who were cured when applying a control given by Figure 3. The patients that converged to the septic death equilibrium (as explained in Section 3.1) are obviously the ones who were not cured with the approach. The criteria to classify successful therapeutic control is to determine if the levels of pathogen (*P*) and damage (*D*) are reduced to very low values (< 0.2). All virtual patients not meeting this criteria were either classified as septic death outcomes if, in addition, the pathogen state did not also approach zero or aseptic death outcomes otherwise. These two latter cases correspond to virtual patients not saved by the applied dosage.

**Fig. 2:**
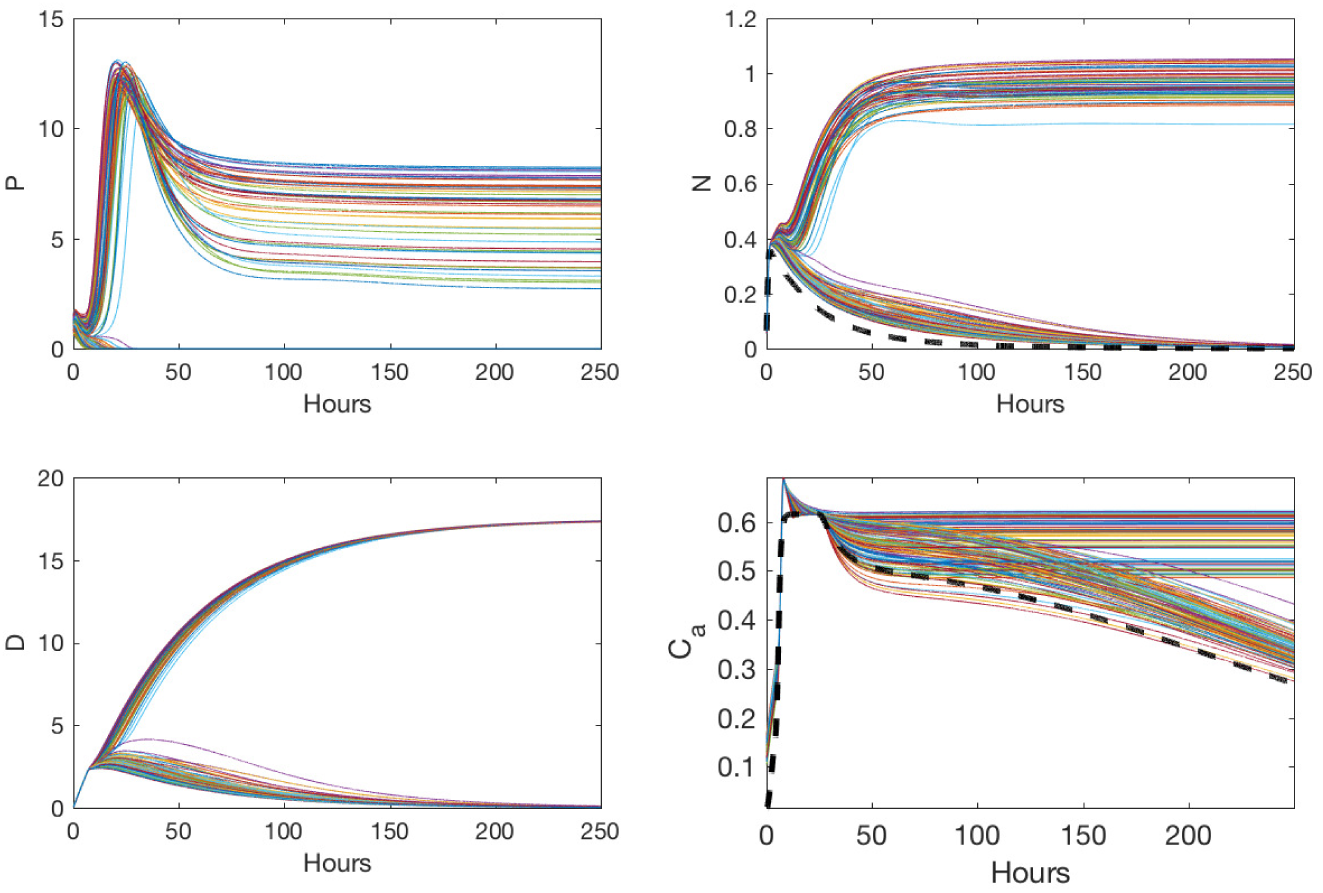
Dashed (- -) curves in the panels for the variables *N* and *C_a_* denote the reference trajectories used in the simulation. The various colored curves display the closed loop state responses for the set of 141 septic patients, of which 92 resolved to the healthy outcome.

**Fig. 3:**
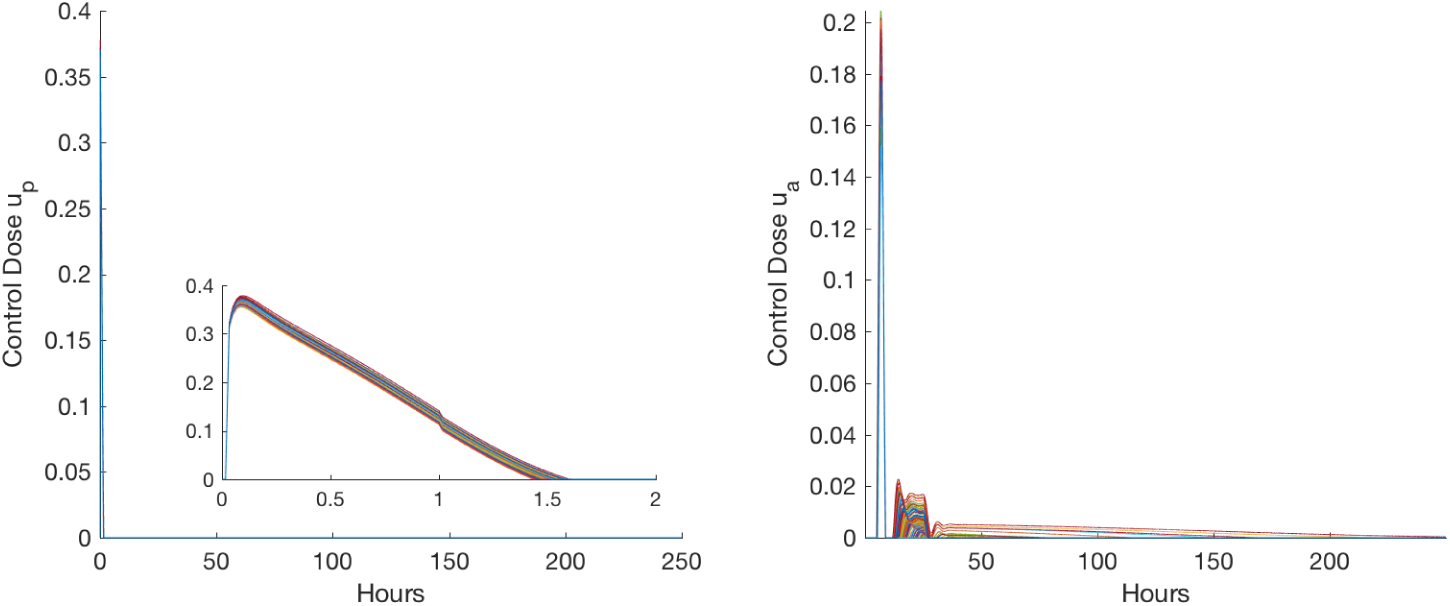
Time evolution of the control *u_p_* and *u_a_* for the set of septic patients on which the strategy was implemented. The zoomed-in plot for *u_p_* provides more details on the duration of the control dose, where the x-axis is shown for only two hours since it is zero for the remaining time

A closer look during the first hours in Figure 3 shows that the amplitude of the control variables is the main difference between different dosing profiles. Similar remarks apply fo *u_a_*. Analyzing *u_a_* shows that it is applied for a longer period of time than *u_p_*, but with smaller amplitude. This was not observed in the optimal control setting of [15], where dosing strategy ended at 30 hrs. It is thus interesting to see if purposefully restraining the dose quantity would have a sizable impact on the result. Surprisingly, however, we observe that forcing the anti-inflammatory control input, *u_a_*, to be zero after 28 hours does not affect the number of cured patients in this current study. This is an important insight to have in order to prevent unnecessary and lengthy dosing protocols. Whereas the maximum duration of the derived optimal control doses in [15] is 30 hours, it is much longer in the model-free control simulations. An extended duration in the model-free setting is the price to pay. On the other hand, the model-free control tracks a single given reference trajectory for all the virtual patients whereas the optimal control strategy [15] strives to infer the trajectories from a mathematical model that is required to be a ‘good’ model for all the virtual patients.

Table 1 displays the results from our study for the 621 patients that qualified for therapy because of sufficiently elevated inflammation. The first column displays the outcomes in the absence of intervention, labeled the *placebo* outcome. Without intervention, 40*%* (252) will resolve to a healthy outcome, while the remaining 60*%* (369) fall into one of the two unhealthy outcome categories. We use the total of 369 unhealthy placebo outcomes to determine the percentage of those that the treatment rescued. Likewise, we use the total of 252 healthy placebo outcomes to determine the percentage of those harmed (i.e. they would have resolved to healthy without treatment but converged to one of the death states instead after receiving treatment). Figures 2 and 3 display the time courses for the sensorless states, P and D. These were guided via the reference trajectories for the states with sensors, *N* and *C_a_*, along with the corresponding control input. The results are reminiscent of [12,14,25]: first apply a large dose of pro-inflammatory therapy, *u_p_*, followed by an anti-inflammatory dose, *u_a_.* The latter attempts to prevent excessive tissue damage resulting from the additional pro-inflammatory signals form the first dose.

**Table 1.** Results of the model-free immune therapy strategy without measurement noise compared to the placebo outcomes.

| Therapy Type: | Placebo | Model-free control therapy |
| --- | --- | --- |
| **Percentage Healthy:** | 40% (252) | 85.66% (518) |
| **Percentage Aseptic:** | 37% (228) | 6.4% (40) |
| **Percentage Septic:** | 23% (141) | 7.8% (49) |
| **Percentage Harmed (out of 252)** | n/a | 0% (0/252) |
| **Percentage Rescued (out of 369)** | n/a | 75.88% (280/369) |

The information we can derive from Table 1 is that the control strategy obviously improves the percentage of cured patients when compared to the placebo case. Our therapy rescued 85.66*%* of the total patient population (621) and 75.88*%* of the combined septic and aseptic population (369). Additionally, 0*%* of the healthy patients are harmed. Figure 4 shows the evolution of the unobservable state *P* and *D* together with the measured states corresponding to *N* and *C_a_* for a set of 228 aseptic patients. Of these, 188 were able to recover from an aseptic placebo outcome, when the generated controls in Figure 5 are applied, driving the pathogen *P* and the level of damage D to zero. Again, one can observe from Figure 4 that some trajectories diverge to the unhealthy aseptic region, where the pathogen is known to have a zero value but the other state variables remain elevated.

**Fig. 4.**
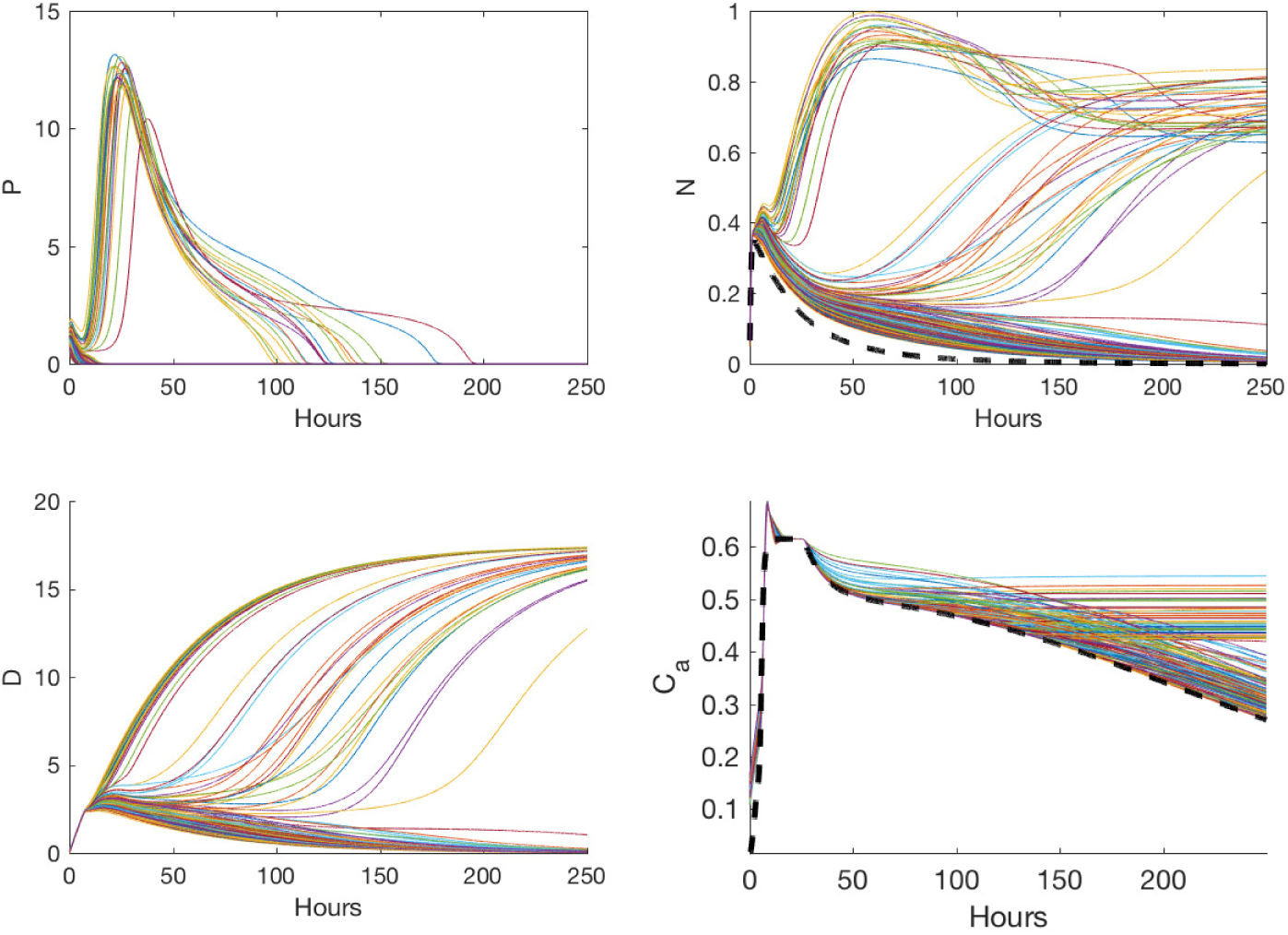
Dashed (- -) curves in the panels for the variables *N* and *C_a_* denote the reference trajectories used in the simulation. The various colored curves display the closed loop state responses results for the set of 228 aseptic placebo patients, of which 188 were cured.

**Fig. 5.**
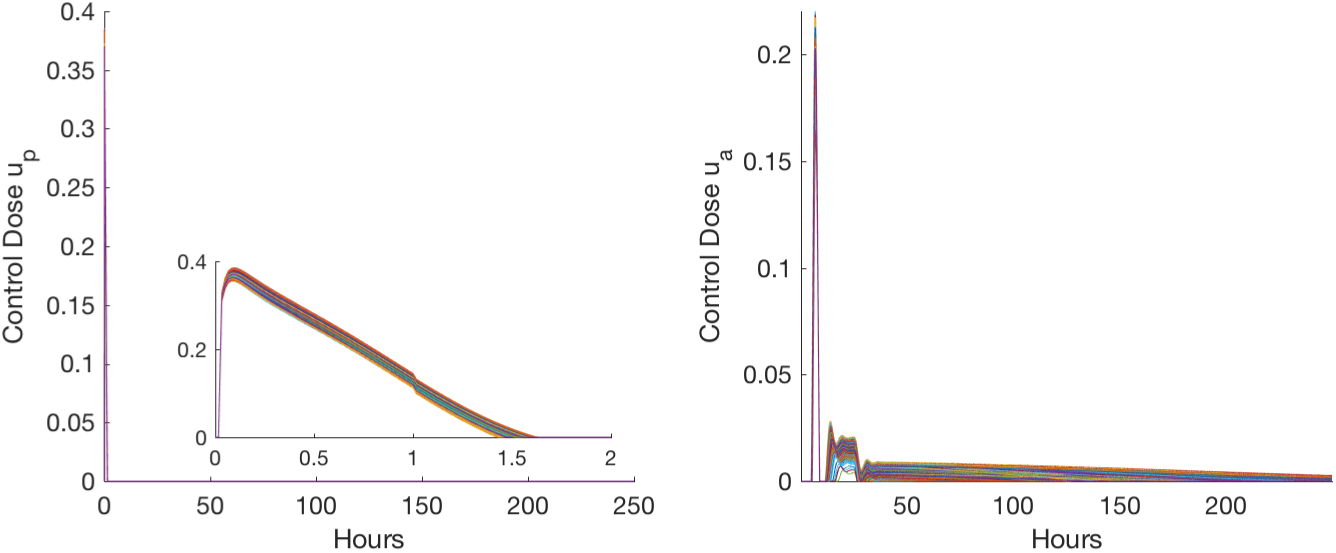
Time evolution of the control inputs *u_p_* and *u_a_* for the set of aseptic patients shown in Figure 4. The zoomed-in plot for *u_p_* provides a better perspective on the duration of the control dose, where the x-axis is shown for two hours only since it is zero afterward.

Overall, the simulation results with respect to successful control of the number of outcomes for both the septic and aseptic placebo outcomes are very encouraging when one considers that only a unique reference trajectory was used for the heterogeneous population. The absence of perfect tracking should not be seen as a weakness of the model free control approach, since the control objective has been attained in most scenarios.

One of the important features of the presented data driven control approach is the necessity to have suitable choice(s) for the reference trajectories. To be more explicit, consider a naive choice of the reference trajectories: a trajectory exponential decaying to zero for *N_ref_* and another trajectory exponentially decaying to the *C_A_* steady state value 0.125. This would not satisfy the control objective since the generated control doses are negative and the level of pathogen will converge to its maximum allowable value. The reason for this behavior can be explained by the fact that the *iP* controller is only concerned about reducing the tracking error without imposing any constraints on the control inputs. Constraints on the control are not implemented simply because the model-free approach is not formulated as an optimization problem. ^2^ However, choosing a reference trajectory that will account for the correct time-varying dynamics of the inflammatory response will eventually generate the correct doses. That is, if we chose, for example, a reference trajectory with a smaller amplitude or with slower rising dynamics, other than what is presented in this work, then it is highly probable that the patient would not converge to the healthy state with respect to the generated control doses received. Similar remarks can be made to Figure 5 as discussed previously for the set of placebo septic patients. It is not surprising to notice a very close pattern with respect to the generated control doses for both sets of septic and aseptic patients. This can be explained in part by a common control objective consisting of tracking the same reference trajectory and also because of what has been discussed before regarding how the inflammatory immune system needs to react in order to eliminate the pathogen without incurring a significant damage.

### 3.4 Results with noise corruption

Consider the effects of corrupting measurement noise on our control problem. Here, a white Gaussian noise is taken into account as in many academic studies, (see, *e.g.*, [19,62]). Otherwise, the same setting as the previous section is kept. Figures 6 and 7 display the states and the corresponding controls for the set of 141 septic placebo patients in which 90 were cured.

**Fig. 6.**
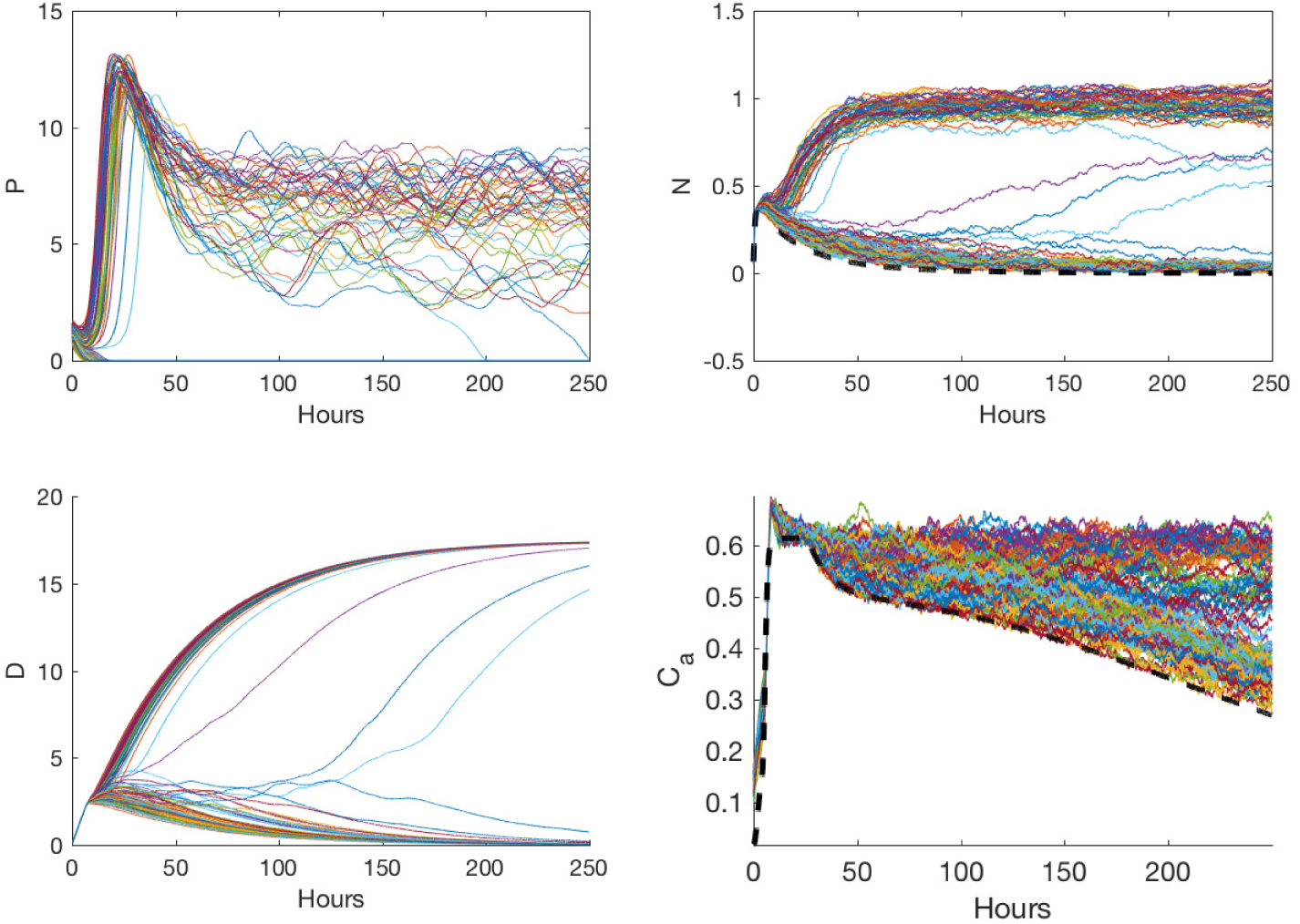
Dashed (- -) curves in the panels for the variables *N* and *C_a_* denote the reference trajectories used in the simulation. The various colored curves display the closed loop state responses for the set of 141 septic placebo patients, of which 90 were cured. Note that the measurements *N* and *C_a_* were corrupted with Gaussian noise.

**Fig. 7.**
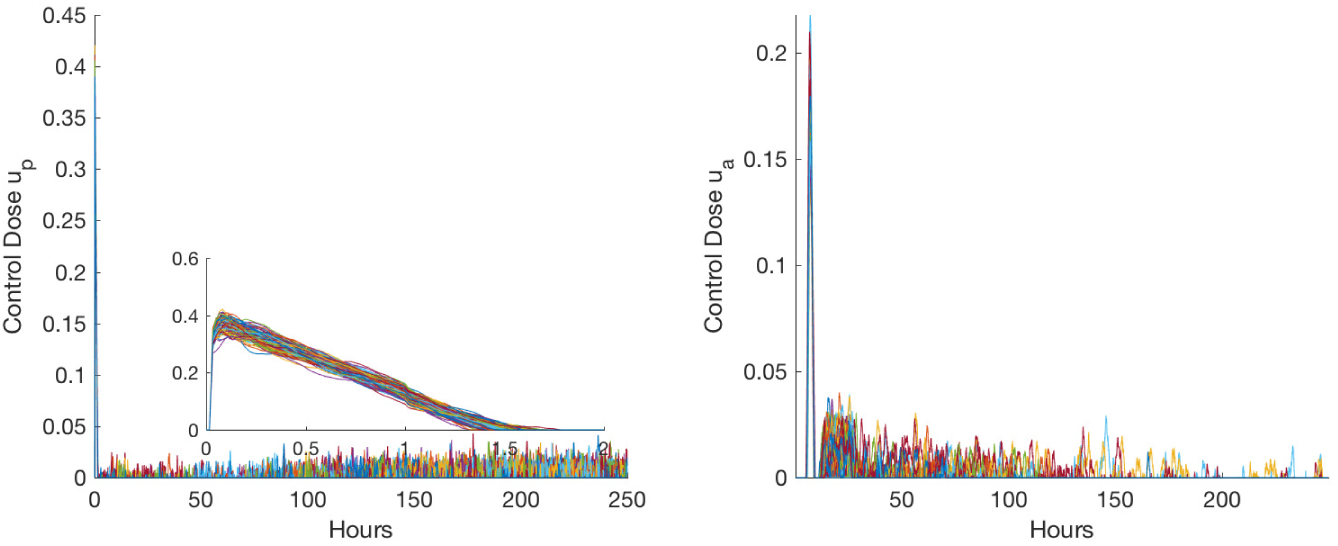
Time evolution of the control *u_p_* and *u_a_* for the set of septic placebo patients when the measurements *N* and *C_a_* were corrupted with Gaussian noise. The zoomed-in plot for *u_p_* provides more detail on the duration of the control dose, with an x-axis shown only for two hours since the doses are zero afterward.

The addition of measurement noise with a standard deviation equal to 10^−3^ only changes the outcome for two of the septic patients, when compared to the initial simulations where no noise was included. However, for the aseptic set of patients, there is a difference of 16 additional patients that did not survive when measurement noise is considered.

*Remark 4* For the model-free simulation with measurement noise, there are mainly two important remarks to make with regard to the discussion of the previous Section. First, for the case of septic patients, restraining the control *u_p_* and *u_a_* to be zero after 2 hours and 28 hours, respectively, will not considerably affect the number of cured patient since 90 patients were cured. One would fail to obtain a similar result when altering the control in the same way for the aseptic case. Although not shown here, a decrease of around 45 patients was observed when compared to the 172 who were cured without restraining control inputs.

## 4 Concluding remarks

In this study we propose a new data-driven control approach in order to appropriately regulate the state of an inflammatory immune response in the presence of pathogenic infection. The performance of the proposed control strategy is investigated in the context of a set of 621 heterogenous model-based virtual patient population having model rate parameter variability. The results of the model-free strategy presented here in the presence of measurement noise are also explored and discussed. The robustness of the approach to parameter variability and noise disturbances is seen in the fact that a single reference trajectory was used to inform the approach about desirable inflmmatory dynamics and from this, the individual dosing strategies found largely produced healthy outcomes. The downside of the proposed control approach to this specific application is the necessity to apply the control for a longer period of time although with small doses. However, we have seen that artificially restricting this small dose from being provided does not affect the outcome of the states in the case when no measurement noise is used; thought it did in the scenarios with measurement noise. We want to emphasize the importance of a suitable choice for the reference trajectory and further studies may provide better insights in this direction.

Past successes of the model-free control feedback approach in other realistic case-studies should certainly be viewed as encouraging for the future development of our approach to the treatment of inflammation. Additionally, the model-free control approach seems to be both theoretically and practically simpler when compared to model-based control designs. This newer viewpoint for control problems in biomedicine needs to be further analyzed in order to confirm its applicability in these complex dynamic systems where the ability to realistically obtain frequent measurement information is limited.

**Fig. 8.**
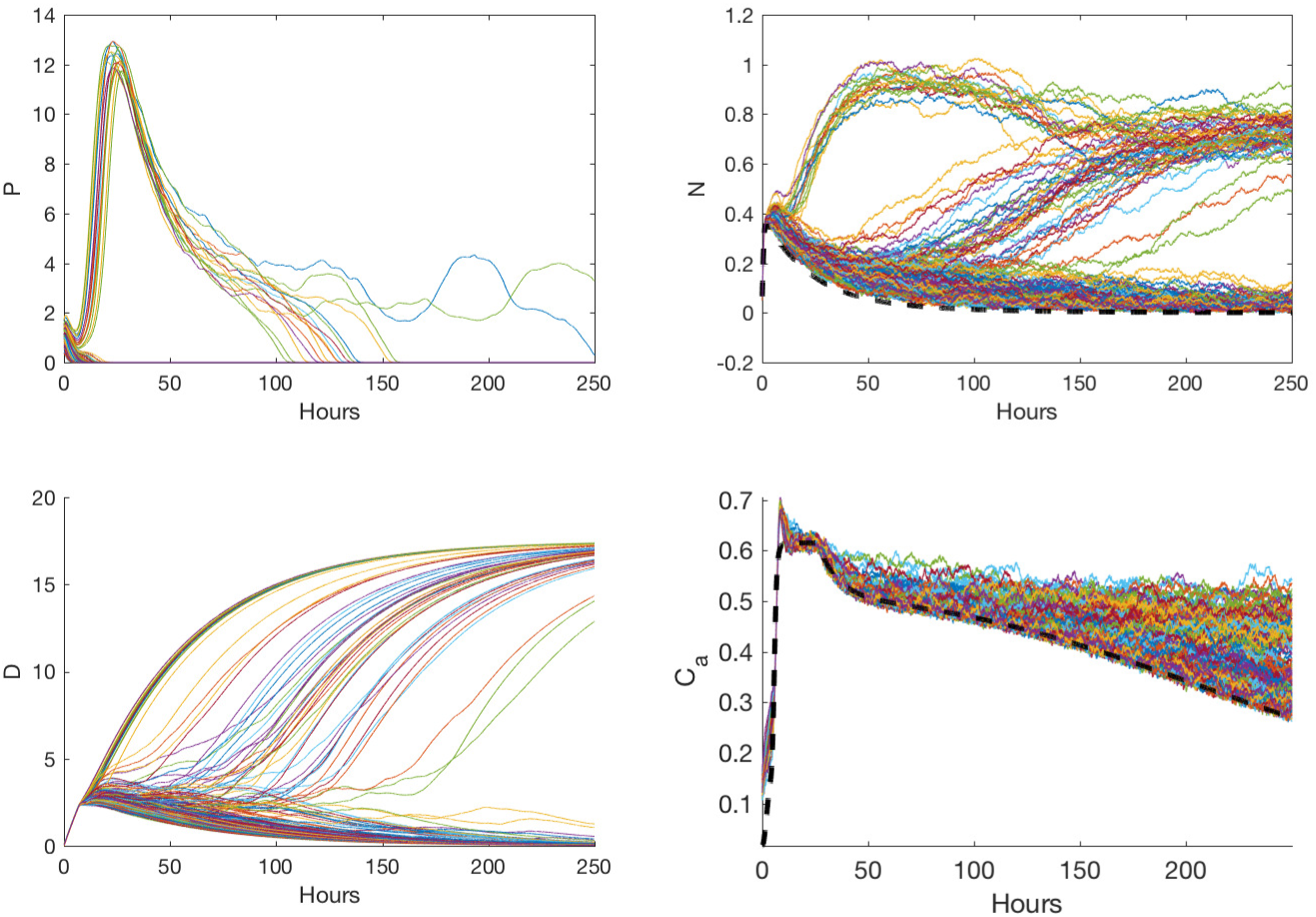
Dashed (- -) curves in the panels showing the time courses of the variables *N* and *C_a_* denote the reference trajectories used in the simulation. The various colored curves display the closed loop state responses for the set of 228 aseptic placebo patients, 172 of which were cured. Note that the measurements *N* and *C_a_* were corrupted with Gaussian noise.

**Fig. 9.**
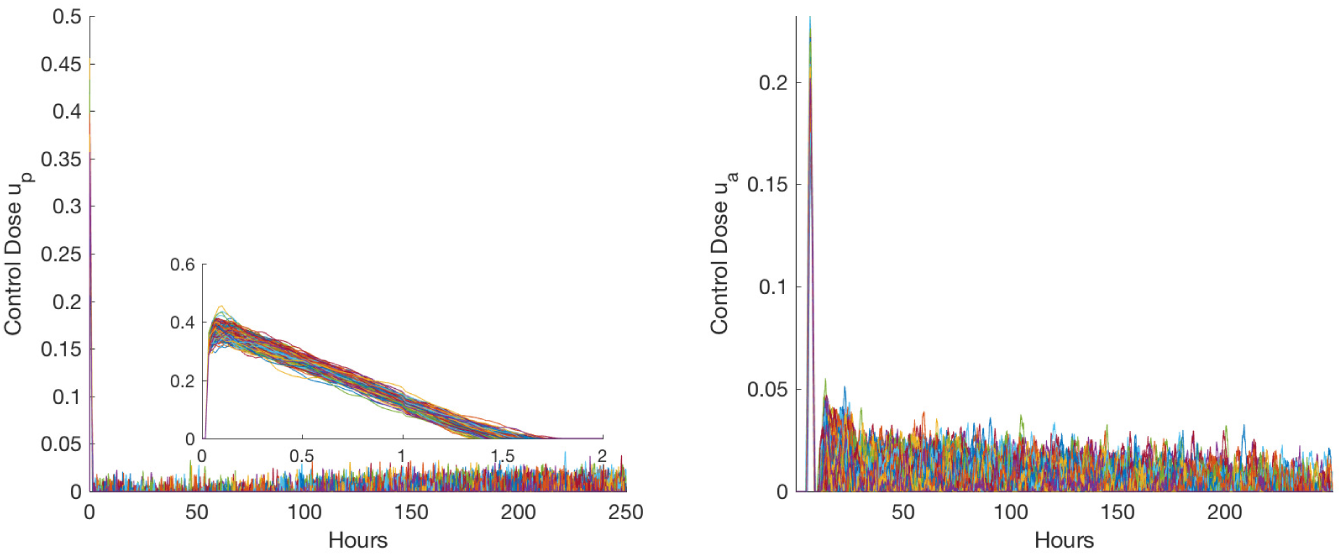
Time evolution of the control *u_p_* and *u_a_* for the set of 228 aseptic placebo patients when the measurements *N* and *C_a_* were corrupted with Gaussian noise. The zoomed-in plot for *u_p_* provides more details on the duration of the control dose, with an x-axis shown only for two hours since the doses are zero afterward.

**Table 2.** Results of the model-free immune therapy strategy with measurement noise compared to the placebo outcomes.

| Therapy Type: | Placebo | Model-free control therapy |
| --- | --- | --- |
| **Percentage Healthy:** | 40% (252) | 82.76% (514) |
| **Percentage Aseptic:** | 37% (228) | 9.02% (56) |
| **Percentage Septic:** | 23% (141) | 8.21% (51) |
| **Percentage Harmed (out of 252)** | n/a | 0% (0/252) |
| **Percentage Rescued (out of 369)** | n/a | 71% (262/369) |

## Footnotes

1 This new viewpoint in control engineering has been successfully illustrated in many concrete casestudies (see, e.g., the references in [34], and [1,2,3,4,17,21,43,47,49,50,52,53,54,57,55,66,72,81,82,87,88,89,90,91,94,95]). Some of the methods have been patented and some have been applied to life sciences [35,43,47,57,81].

2 Allowing the control to be only positive semidefinite will result in a zero control all the time.

